# Mapping of the viral shunt across widespread coccolithophore blooms using metabolic biomarkers

**DOI:** 10.1101/2024.11.03.621239

**Authors:** Constanze Kuhlisch, Guy Schleyer, J. Michel Flores, Flora Vincent, Marine Vallet, Daniella Schatz, Assaf Vardi

**Affiliations:** Department of Plant and Environmental Sciences, Weizmann Institute of Science, 7610001 Rehovot, Israel; Department of Marine Microbiology and Biogeochemistry, NIOZ Royal Netherlands Institute for Sea Research, 1797 SZ Texel, Netherlands; Department of Earth and Planetary Sciences, Weizmann Institute of Science, 7610001 Rehovot, Israel; Developmental Biology Unit, European Molecular Biological Laboratory, 69117 Heidelberg, Germany; Max Planck Fellow Group Plankton Community Interaction, Max Planck Institute for Chemical Ecology, 07745 Jena, Germany; Institute for Inorganic and Analytical Chemistry, Friedrich Schiller University Jena, 07743 Jena, Germany

**Keywords:** algal bloom, *Gephyrocapsa huxleyi*, alga-virus interaction, virocell, vDOM, halogenation, biomarker

## Abstract

The viral shunt is a fundamental ecosystem process which diverts the flux of organic carbon from grazers to heterotrophic microorganisms. Through the extracellular release of metabolites, lytic viral infections supply 2-10% of photosynthetically fixed carbon in the ocean for bacterial respiration. Despite its significance for the carbon cycle, we lack tools to detect the viral shunt in the natural environment and assess its ecological impact. Here, we study the use of exometabolites as biomarkers for the viral shunt by applying molecular, metabolomics, and oceanographic tools in blooms of the cosmopolitan microalga *Gephyrocapsa huxleyi* across the Atlantic Ocean, spanning four biogeochemical provinces between Iceland and Patagonia. We mapped the distinct metabolic footprint of its viral infections using exo- and endometabolomics approaches and detected nineteen organohalogen metabolites across the blooms, showing their global formation. Time-resolved comparison of particulate and dissolved metabolite pools during an induced mesocosm bloom indicated virocells – actively infected host cells – as the source of the halogenated metabolites. Three trichloro-iodo metabolites were present during demise of all virus-infected blooms, highlighting them as suitable metabolic biomarkers. The environmental stability of these halometabolites in the DOM pool over a few days can recapitulate viral infections at earlier stages of phytoplankton bloom succession.

## Introduction

Viruses are the most abundant biological entities in the ocean with estimated 10^7^ viral particles per mL seawater (1). As infective agents, viruses are key players in the marine environment that can infect between 1% and 60% of their host populations (2, 3). During interaction with their hosts, viruses remodel host cell metabolism and introduce auxiliary metabolic genes, thus creating a distinct cell state referred to as virocell (4, 5). The virocell metabolism has important consequences for the chemical composition of the surrounding seawater (6-8). The release of all intracellular metabolites into the dissolved organic matter (DOM) pool at the end of a lytic infection diverts the carbon flux away from grazers to heterotrophic microorganisms such as bacteria (9). This fundamental ecosystem pathway known as the ‘viral shunt’ enhances microbial respiration in the surface ocean and impacts the fate of carbon in microbial food webs. However, we still lack tools to map the carbon flux through the viral shunt and thereby assess the impact of viral lysis on the marine carbon cycle. Recently, new methods have been developed to quantify ongoing viral infections at the single-cell level (3, 10, 11) and taxon-specific cell lysis rates via extracellular ribosomal RNA (12) in the marine environment. The composition of virus-induced DOM (vDOM) differs from exudates and lysates of non-infected cells (7), suggesting that taxon-specific vDOM signatures can be used as metabolic biomarkers to quantify the viral shunt of specific algal blooms in the marine environment.

The cosmopolitan bloom-forming microalga *Gephyrocapsa huxleyi* (formerly known as *Emiliania huxleyi*) is a key player in the marine carbon and sulfur cycles, and a sensitive reporter of the ecological consequences of climate change, such as ocean acidification and poleward species expansion (13-15). Viral infection by the *Emiliania huxleyi* virus (EhV) is a major cause of mortality in the large-scale blooms of *G. huxleyi*, controlling the metabolic landscape during bloom demise. In a recent mesocosm experiment, we induced *G. huxleyi* blooms and followed changes in the exometabolic landscape throughout bloom succession (6). Viral infection led to a distinct metabolic footprint during bloom demise characterized by metabolites with a dual halogenation of chlorine and iodine that are rare in the marine environment (6). Despite the formation of volatile organohalogens that contribute to atmospheric ozone depletion (16), little is known about the role of marine phytoplankton blooms as a source for halogenated metabolites, which are characterized by their high bioactivities and environmental persistence. Since there are limited diagnostic tools to map the viral shunt in the ocean, we aimed to assess the use of the dual halogenated metabolites as possible biomarkers for the viral shunt in widespread blooms of *G. huxleyi* across the Atlantic Ocean.

## Results

### Sampling oceanic blooms at various phases

To encompass the wide distribution range of *G. huxleyi* and its association with temperate and subpolar waters (17), several phytoplankton blooms were sampled across the Atlantic Ocean, spanning four biogeochemical provinces between Iceland and Patagonia (18) (**Figure 1A, Dataset S1**). The sampling included three Boreal summer blooms in the Northeast Atlantic (V1-2, B1), four Austral summer blooms in the Southwest Atlantic (G1-4), and two non-bloom reference sites (G5-6). Four of these blooms were followed in Lagrangian drifts which allowed the repeated sampling of these blooms for 3-5 days (V2, B1, G3, G5). Coccolithophores have a distinct optical signature based on the scattering of light from their calcite exoskeleton (**Figure 1B**) that can be detected by satellite remote sensing. *G. huxleyi* blooms were thus identified using satellite imagery taken by the MODIS/Aqua instrument, with blooms being characterized by an elevated particulate inorganic carbon (PIC) signal of >0.7 mmol m^-3^ (19). PIC ranged from as low as 0.3 mmol m^-3^ (Gayoso G1) to ∼2 mmol m^-3^ (VICE V1 and V2, Gayoso G3 and G4) and as high as 2.5 mmol m^-3^ (Breizh B1; **Dataset S1**). In addition, the *in*-*situ* abundance of *G. huxleyi* was enumerated using on-board flow cytometry. The calcite exoskeleton of *G. huxleyi* increases the side scattering of light and distinguishes its cells from other nanoeukaryotic phytoplankton. Blooms in the open ocean typically reach cell densities of 1-10×10^6^ cells L^-1^ (17). In correspondence with the satellite PIC signal, *G. huxleyi* cells were present at all locations, with cell abundances reaching as high as 3.7×10^6^ cells L^-1^, indicating intense blooms (B1 and G4; **Figure 1C**). A moderate PIC signal, despite low abundances of <1×10^6^ cells L^-1^ (V1 and V2), may be explained by light scattering of coccoliths that were shed from dying cells during bloom demise, while the low PIC signal at G1, despite a high cell abundance, is due to the deep bloom depth of ∼50 m. The temporal bloom dynamics at each location were assessed by mapping satellite data of PIC and chlorophyll *a* (Chl *a*) around the time of sampling (**Figure S1**, see S1B for a detailed example). Pre- and post-bloom phases are defined by having <0.7 mmol m^-3^ PIC. During the bloom phase, the PIC signal increases with the peak of the bloom and onset of the demise phase being marked by reaching a minimum Chl *a*:PIC ratio (20). During the demise phase, the Chl *a*:PIC ratio increases due to the attenuation of the PIC signal. The Boreal blooms V1 and V2 were in an early and late demise phase, respectively. The five-day drift sampling (B1) captured the onset of demise of an intense bloom of *G. huxleyi* (>3×10^6^ cells L^-1^). The Austral bloom samplings captured a bloom phase (G4), an early demise phase (G3) and two late demise phases (G1, G2). The sampling of several *G. huxleyi* blooms in their demise phase allowed to examine whether this was the result of viral infection and whether metabolic biomarkers can be applied to report the formation of DOM through the viral shunt.

**Figure 1.**
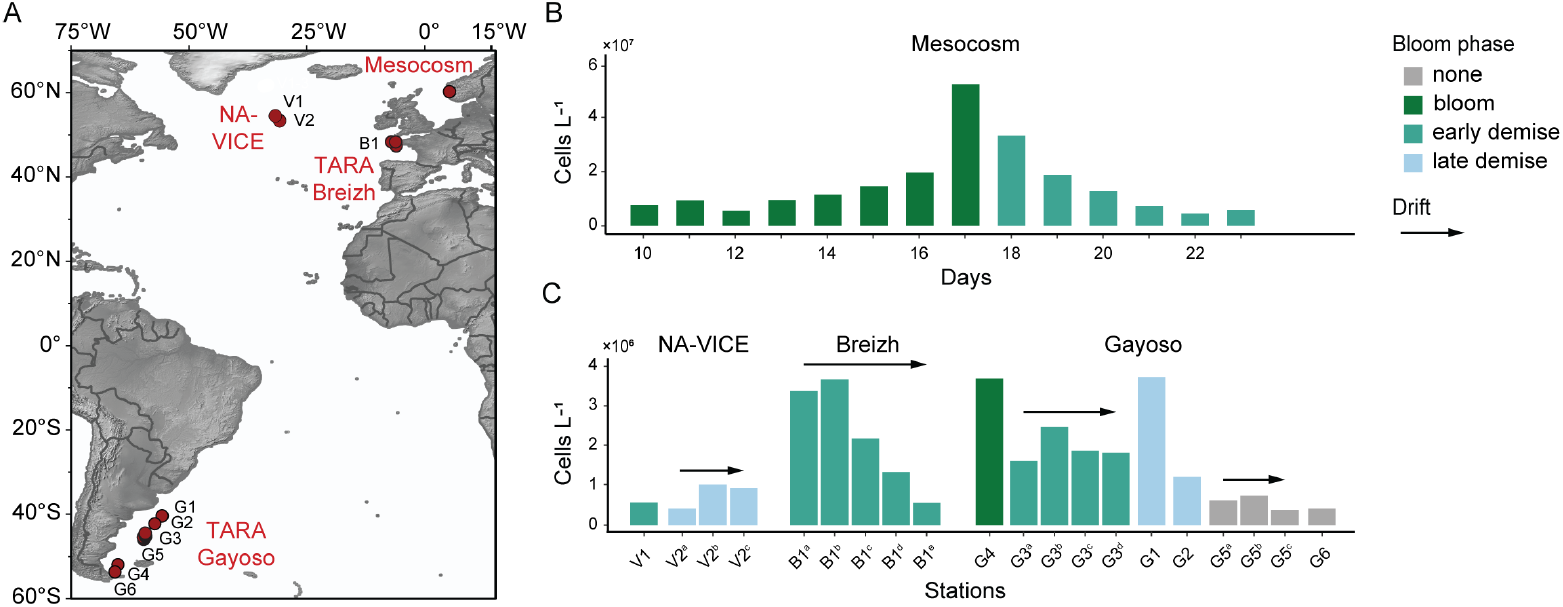
*G. huxleyi* blooms across the Atlantic sampled in various bloom phases. A) Map shows sampling locations of *G. huxleyi* blooms including an induced coastal mesocosm bloom, oceanic Boreal summer blooms (NA-VICE cruise, TARA Breizh cruise), and oceanic Austral summer blooms (TARA Gayoso cruise). B) At the peak of the induced mesocosm bloom, calcified *G. huxleyi* cell abundance reached over 10^7^ cells L^-1^. C) *In*-*situ* abundances of calcified *G. huxleyi* cells in the oceanic blooms were tenfold lower. Counts depict the maximum abundance reached in each depth profile. The bar color indicates the bloom phase at the time of sampling based on the temporal dynamics of flow cytometry counts (mesocosm bloom) and satellite images (oceanic blooms; see Figure S1B for details). Arrows indicate Lagrangian drifts with a consecutive sampling of the same water mass.

### Metabolic footprints of viral infection in bloom demise

Oceanic bloom samplings often provide snapshots in time and require molecular and metabolic biomarkers to assess the bloom and infection states. The vDOM of induced *G. huxleyi* blooms is characterized by several novel metabolites with a dual halogenation of chlorine and iodine (6) (**Figure 2A**). This unique metabolic hallmark renders them potential metabolic biomarkers for the viral shunt in oceanic *G. huxleyi* blooms. To assess whether this dual halogenation differs in its occurrence from other halogen-bearing exometabolites, we mapped all halogenated metabolites throughout the phytoplankton bloom succession of the induced mesocosm blooms (**Figure S2A**). Iodinated metabolites were detected specifically during the *G. huxleyi* bloom demise and depended on the extent of viral infection. In contrast, brominated and chlorinated metabolites were already present extracellularly at the onset of the *G. huxleyi* bloom and increased during the demise phase independent of the infection level. The total concentration of chloro-iodo metabolites was strongly correlated with viral abundances (cell-bound and free EhV; ρ = 0.82-0.83, p < 0.01; **Figure 2B**). Furthermore, a weak correlation was found with virus-derived glycosphingolipids (vGSLs; ρ = 0.65, p = 0.11) that serve as intracellular biomarkers for viral infection. No positive correlation was found with the abundance of *G. huxleyi* and other phytoplankton community members, such as nano- and picoeukaryotes, bacteria, or physicochemical water properties (**Figure S2B**). These observations reinforce the link between the formation of the chloro-iodo metabolites and active viral infection, and highlights the possibility to utilize iodinated metabolites, specifically chloro-iodo metabolites, as metabolic biomarkers for vDOM formation in open ocean blooms of *G. huxleyi*. The high correlation (ρ = 0.97, p < 0.01) between the two intracellular infection markers (cell-bound EhV and vGSLs; **Figure S2B**) emphasizes that intra- and extracellular viral infection markers have different dynamics: intracellular markers report the virocell state, while extracellular chloro-iodo metabolites report the post-lysis state of vDOM formation.

**Figure 2.**
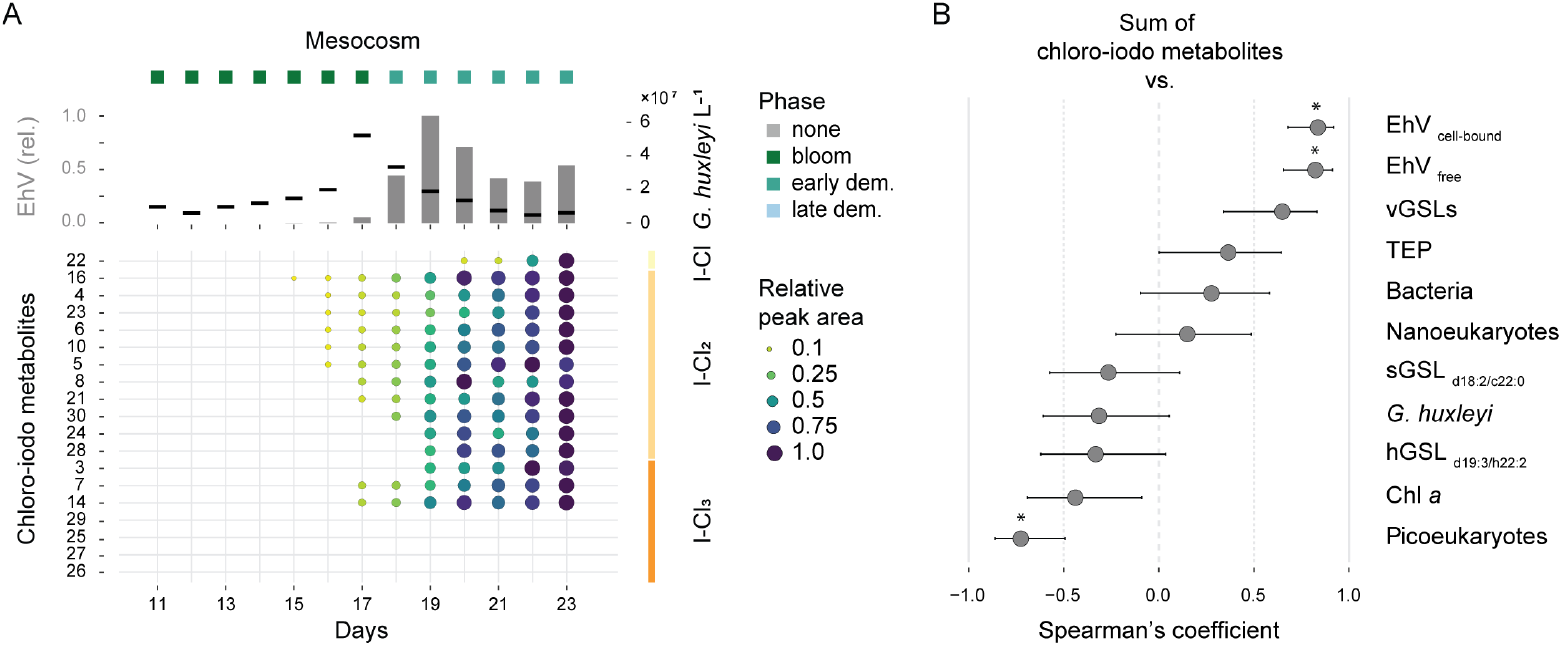
Chloro-iodo metabolites correlate with viral infection markers in induced *G. huxleyi* blooms. A) Occurrence of chloro-iodo exometabolites (bottom panel) is strongly linked to viral infection (top panel) during the demise of an induced mesocosm bloom. Abundance of *G. huxleyi* cells and intracellular viruses (top panel) delineates the bloom and infection dynamics. Abundance of EhV (*E. huxleyi* virus) is scaled to the maximum abundance. Bubble size and color represent relative peak area (normalized and scaled per metabolite). Bloom phases and Lagrangian drifts are indicated. B) Spearman’s coefficients and 95% CIs for the correlation between the abundance of total chloro-iodo exometabolites and various bloom and demise descriptors for induced mesocosm blooms. Variables include the abundance of cell-bound and free EhV particles, lipid markers of virus-infected (vGSL), susceptible (sGSL) and all *G. huxleyi* cells (hGSL), cell abundance of *G. huxleyi*, nanoeukaryotes, picoeukaryotes and bacteria, total Chl *a*, and transparent exopolymer particles (TEP). Stars (*) indicate significant correlations (p < 0.05).

We aimed to examine the sensitivity of the chloro-iodo metabolites as metabolic biomarkers in the natural environment. Using a liquid chromatography-mass spectrometry (LC-MS)-based metabolomics approach, we profiled diverse oceanic blooms covering the widespread occurrence of *G. huxleyi*, from Iceland to Patagonia, and different bloom phases. Despite the tenfold lower cell abundances compared to induced mesocosm blooms, we detected several chloro-iodo metabolites in the oceanic *G. huxleyi* blooms (**Figure 3**). In total, a suite of 19 chloro-iodo metabolites was detected intra- and extracellularly in the environmental samples by applying an untargeted metabolite profiling approach using LC-MS in positive and negative ionization modes. Most detected metabolites were dichlorinated (11 metabolites) and trichlorinated (7 metabolites), with one metabolite being monochlorinated. The largest number (9-11 metabolites) was detected in filter samples from two North Atlantic blooms in their demise phase (V1, V2). *G. huxleyi* abundances were low in both locations (<10^6^ cells L^-1^), while the infection level was higher at the early demise phase (V1), as reflected by intracellular EhV abundances enumerated by qPCR of the major capsid protein (*mcp*) gene. Chloro-iodo metabolites were found with higher intensities at the early demise phase (V1), and the weak metabolic signature at the late demise phase further dissipated during the 3-day drift (V2a-c). During the drift with the intense Boreal summer bloom (B1a-d), four chloro-iodo metabolites were detected at similar intensities throughout the drift period. Satellite imagery showed that this bloom reached the lowest Chl *a*:PIC ratio during the drift period compared to the preceding and following week (**Figure S1**). *In*-*situ* measurements showed decreasing cell abundances and slightly increasing intracellular EhV abundances, suggesting it to be virus-infected bloom at the onset of the demise phase. Among the Austral summer blooms, the highest number and intensity of chloro-iodo metabolites were found in a demised bloom (G2) that showed the highest EhV abundance along the Patagonian Shelf. In an adjacent post-bloom phase station (G1), two metabolites were detected at low intensity. No chloro-iodo metabolites were found at the bloom phase stations (G4, G5) that showed higher cell counts (1.6-3.7×10^6^ cells L^-1^) but lower EhV abundances, and at the reference sites (G3, G6). Notably, there is a difference in the occurrence pattern of intracellular viruses – reporting a pre-lysis state – and of extracellular chloro-iodo metabolites – reporting a post-lysis state. For example, while the intracellular viral abundance at the late demise phase station G2 was low, several chloro-iodo metabolites were detected extracellularly. This may indicate the difference in the fate of intracellular biomarkers, e.g. by cell lysis and sedimentation, compared to dissolved metabolites, e.g. by bacterial degradation and lateral transport via ocean currents. The detection of several chloro-iodo metabolites in their demise phase in oceanic blooms suggests that they can be used as sensitive metabolic biomarkers to assess the impact of lytic viral infection through the viral shunt on the biogeochemistry and succession of *G. huxleyi* blooms.

**Figure 3.**
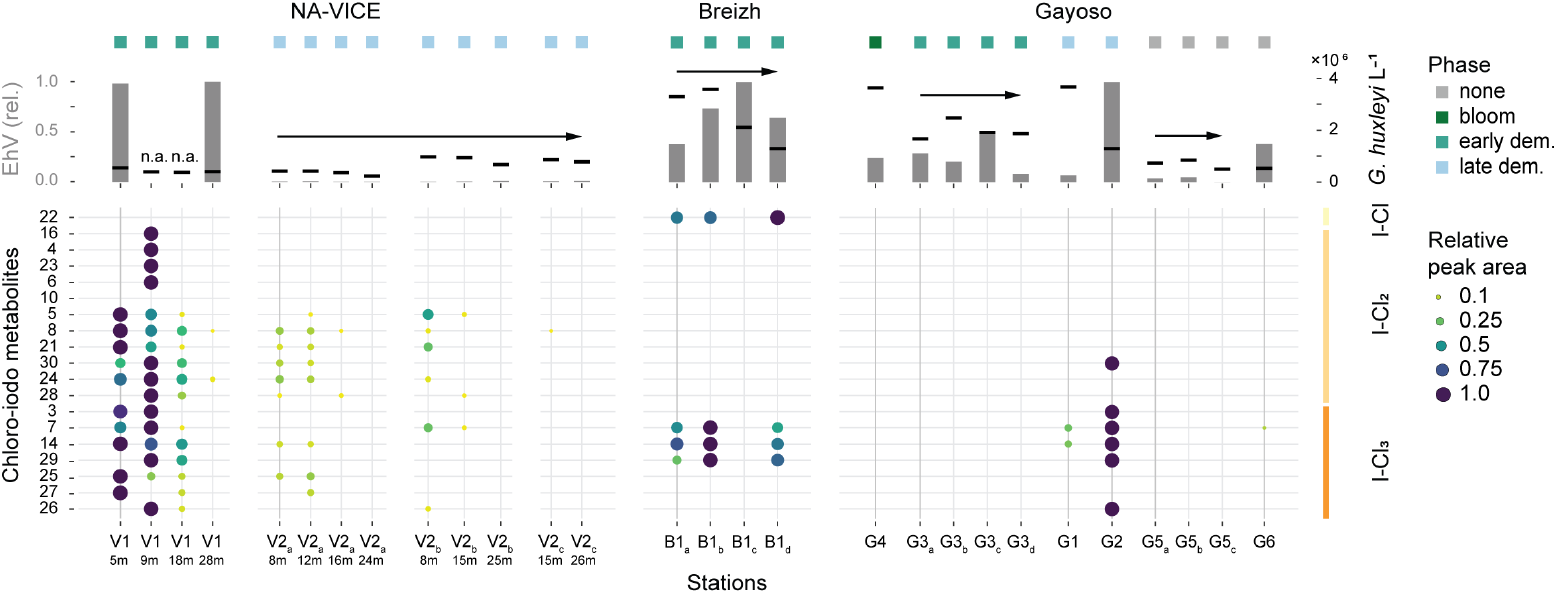
Occurrence of chloro-iodo metabolites in infected oceanic *G. huxleyi* blooms at demise and post-bloom phases. Chloro-iodo metabolites occur in oceanic blooms of *G. huxleyi* at various demise phases (bottom panel). Abundances of *G. huxleyi* and intracellular EhV (top panel) delineates the bloom and infection states (EhV scaled per cruise). Bubble size and color represent relative peak area (normalized and scaled per metabolite). Bloom phases and Lagrangian drifts are indicated. n.a. – not available.

### Tracking the transition from intra-to extracellular carbon pools through viral lysis

To assess the transfer of the chloro-iodo metabolites to the marine DOM and possible transformations by biotic and abiotic processes, we compared their presence within the DOM fraction, representing the extracellular pool and the particulate organic matter (POM) fraction, representing the intracellular pool. The high temporal resolution of the metabolic profiling of the induced blooms enabled us to map the chloro-iodo metabolites over 11 days and identify variations in both pools (**Figure 4**). Two metabolites (#25 and #27) were found solely as cellular metabolites (vPOM) across all blooms and may have rapid turnover times upon cell lysis, e.g., by bacterial degradation or chemical transformation, making them part of the highly labile DOM. Six chloro-iodo metabolites were found solely extracellularly. Of these, metabolite #22 was present exclusively as vDOM across all bloom samples (**Dataset S2**), suggesting an extracellular origin, e.g., as a transformation product of the viral lysate by the heterotrophic microbial community. About half of the chloro-iodo metabolites (9 metabolites) were found intrac- and extracellularly, suggesting a cellular origin and flux into the DOM. The intracellular origin is further supported by the observation that most of these metabolites were first detected in the POM fraction. However, they were enriched extracellularly by the end of the demise phase. The accumulation of chloro-iodo metabolites in the DOM suggests that they are stable for at least a few days. This finding is supported by the reoccurring detection of chloro-iodo metabolites during the drift with a demising bloom in the North Atlantic (**Figure 2**) and suggests that some of these metabolites may be recalcitrant to bacterial degradation. The release and stability of the chloro-iodo metabolites in the DOM pool can capture and recapitulate the dynamics of viral infections at earlier stages of the bloom dynamics.

**Figure 4.**
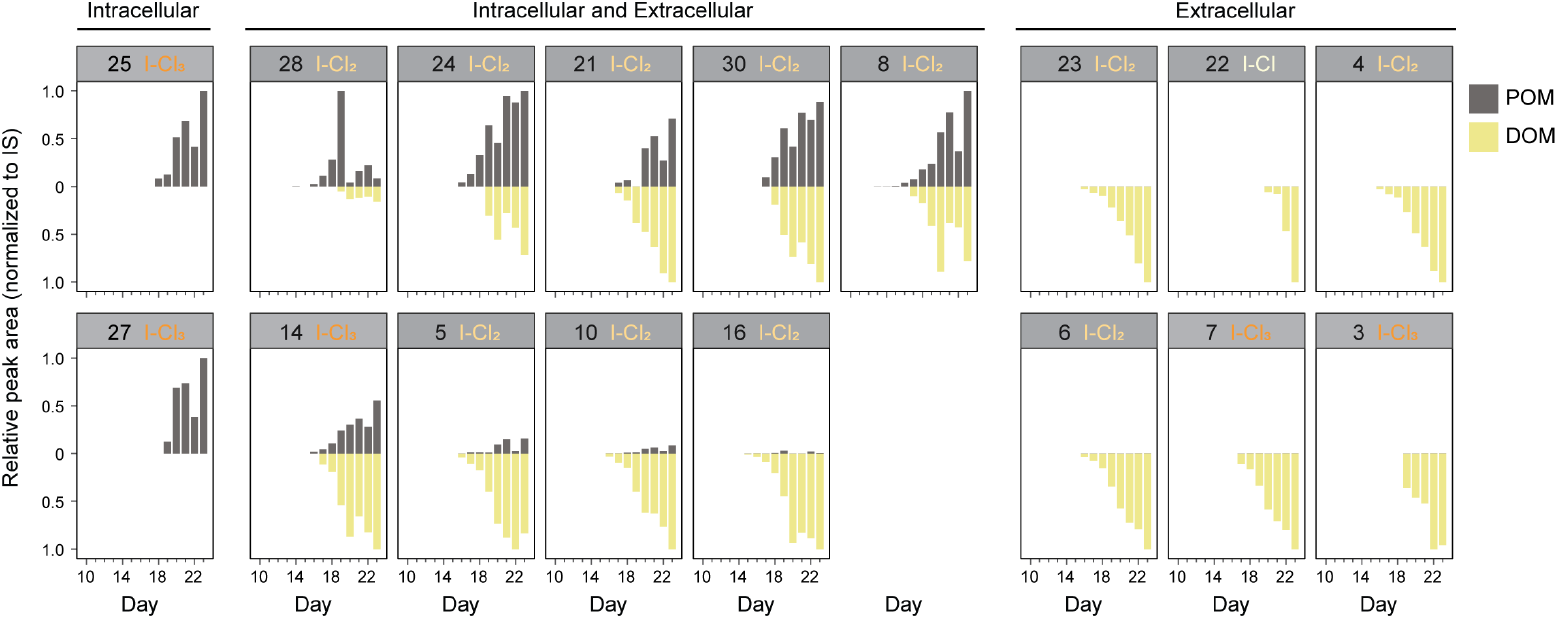
Detection of chloro-iodo metabolites in intra- and extracellular organic matter pools. Profiling the temporal dynamics of chloro-iodo metabolites during host-virus interaction in a mesocosm bloom of *G. huxleyi*. Metabolites were quantified from filter samples (POM, intracellular; black) and filtrates (DOM, extracellular; yellow) of a virus-infected mesocosm bloom. Peak areas were normalized to the internal standard (IS) and scaled to the highest abundance of each metabolite. Metabolites are grouped according to their detection in these pools – only in the intracellular pool, only in the extracellular pool, or both pools – and further ordered by the intra-to-extracellular ratio.

### Trichloro-iodo metabolites are sensitive metabolic biomarkers

Metabolites #7, #14 and #29 were detected frequently in samples from the North and South Atlantic (**Figure 5A**), suggesting that they are most sensitive as metabolic biomarkers in the natural environment. They were detected both at early (V1, B1) and late (V2, G2) demise phases. Interestingly, in the demise phase of an intense bloom in the North Atlantic (B1), metabolites #7 and #14 were detected extracellularly and, at lower and decreasing signal intensities, intracellularly (**Figure 5B**), while vGSLs as intracellular infection markers were not detected (**Figure S3A**). This suggests that chloro-iodo metabolites can report viral infections of past bloom events even when intracellular infection markers, such as vGSLs and *mcp* (viral major capsid protein gene), are low or already diminished below detection limit following the cell lysis at the end of the infection. All three halometabolites are trichloro-iodo metabolites, and metabolite #7 seems to be a hydroxylated derivative of #14 based on predicted elemental composition and similar mass fragmentation patterns (**Dataset 2, Figure S3B**). While metabolites #7 and #14 both increased in the DOM pool throughout the demise of the induced mesocosm bloom, they showed a different partitioning between the POM and DOM pools, with a higher prevalence of metabolite #7 in the DOM and a higher prevalence of metabolite #14 in the POM (**Figure 4**). As metabolite #29 was found in the oceanic blooms with a similar occurrence pattern to #7 and #14, it may function as an additional metabolic biomarker despite its lower signal intensities (**Figure 5A**). The absence of metabolite #29 in the mesocosm bloom suggests site-specific metabolic variability governed by e.g., host and virus strain differences and metabolic plasticity. As the detection of metabolites #7 and #14 in environmental samples appeared robust regarding various extraction and analysis approaches (**Figure S3C**), they can be easily incorporated into environmental sampling strategies.

**Figure 5.**
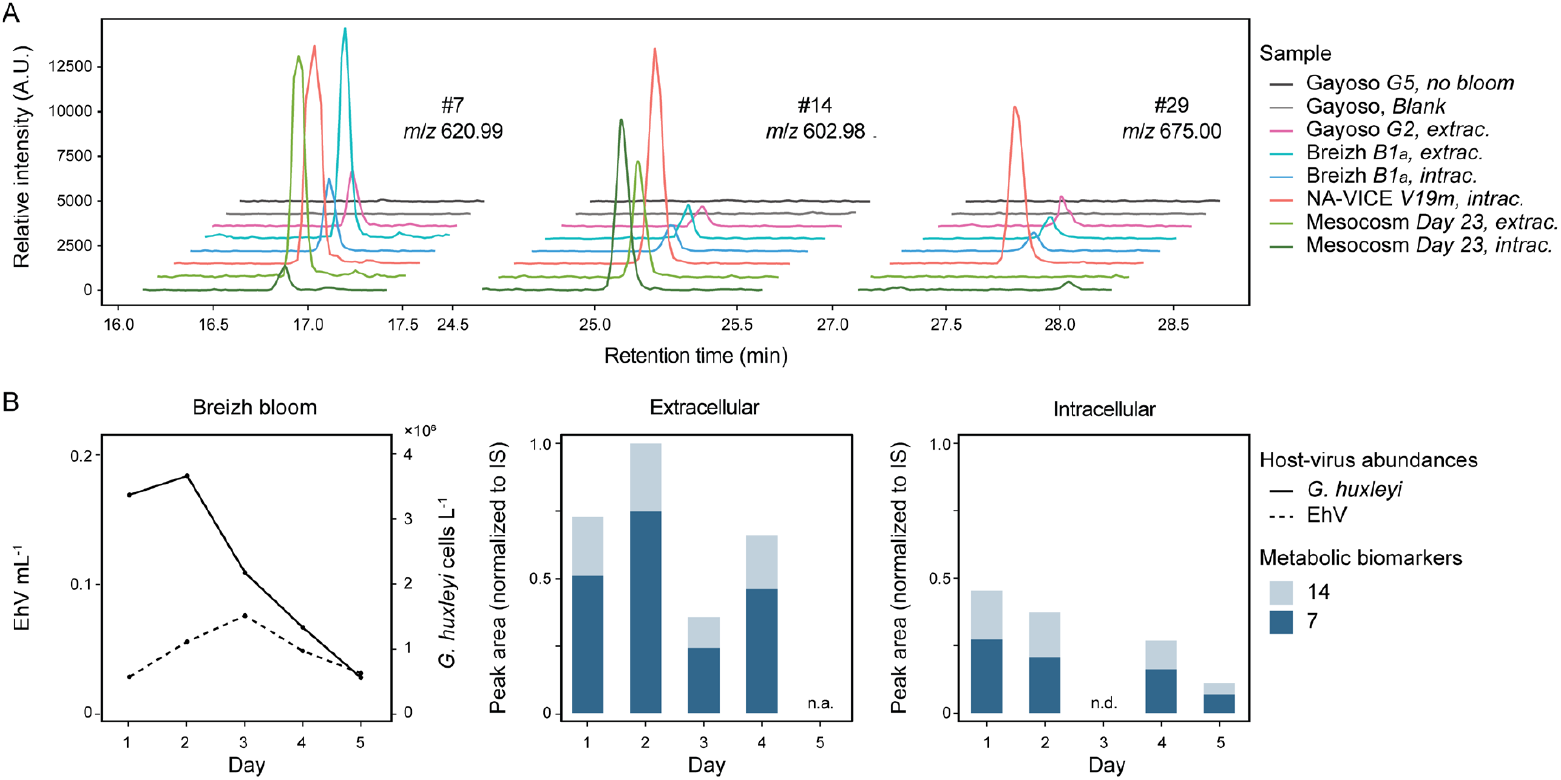
Trichloro-iodo metabolites are widespread and sensitive metabolic biomarkers for the viral shunt in natural *G. huxleyi* blooms. A) Extracted ion chromatograms of the [M-H]^-^ adducts show the sensitive detection of metabolites #7, #14 and #29 in representative samples from oceanic and induced blooms. Metabolites were absent in non-bloom and artificial seawater blank samples. B) During the Lagrangian drift with a virus-infected, demising North Atlantic bloom (B1), decreasing cell abundances of *G. huxleyi* (solid line) and low levels of intracellular EhV (dotted line) were found. Chloro-iodo metabolites #7 and #14 were detected with a decreasing trend extracellularly and at lower intensities intracellularly (peak areas were normalized to the internal standard (IS) and scaled to Day 2). n.a. – not available. n.d. – not detected.

## Discussion

Molecular biomarkers are powerful tools to assess host-pathogen interactions such as viral infections in the marine environment. For *G. huxleyi*, several gene-, transcript- and lipid-based biomarkers have been identified that can shed light on viral infections of natural blooms. The viral gene marker *mcp* allows to assess the abundance of EhV virions intra- and extracellularly (20), while gene expression analyses can resolve the fraction of actively infected cells (21) and the various states of the infection cycle (11, 22). In addition, a suite of lipid markers can reveal the presence of susceptible (sGSLs), infected (vGSLs), and resistant (resistance-specific GSLs, rsGSLs) cells in *G. huxleyi* blooms (23, 24). While this suite of molecular biomarkers allows to characterize viral infections at the cellular level, we lack biomarkers to study infection outcomes beyond host cell death, such as the carbon flux into the DOM pool through the viral shunt. So far, the viral shunt has been assessed only indirectly by measuring host and virus abundances and their specific growth rates (25). The chloro-iodo metabolites assessed in this study are promising candidates as metabolic biomarkers for directly quantifying the formation of vDOM in *G. huxleyi* blooms, due to their specificity, sensitivity, and environmental stability.

Both abiotic and biotic processes can lead to the formation of halogenated metabolites in the marine DOM pool. The chloro-iodo metabolites of the *G. huxleyi* vDOM are aliphatic metabolites, as suggested by their elemental composition and H/C ratio (**Figure S4, Dataset S2**). Aliphatic substrates are mainly halogenated via radical enzymatic pathways mediated by Fe(II)/α-ketoglutarate-dependent halogenases and a few Vanadium- and flavin-dependent halogenases (26-28). The presence of 2-3 double bonds further suggests the involvement of halogenases that employ an electrophilic mechanism targeting electron-rich substrates via alkene addition or aromatic substitution reactions. However, we currently lack knowledge on the presence and role of halogenases in blooms of *G. huxleyi*. In addition to enzymatic halogenations, sunlight can also lead to organohalogen formation (29). Photosensitized DOM and reactive oxygen species can induce reactive halide species that non-specifically halogenate metabolites. The seawater concentration of bromide, photolability of iodinated DOM (30), and reduction potential of chloride (31) lead to a preferential bromination, however, no brominated exometabolites were found as constituents of the *G. huxleyi* vDOM. Furthermore, photoiodination preferentially affects lignin-like substrates (32), strengthening the role of enzymatic pathways in forming the aliphatic chloro-iodo metabolites. Strikingly, other aliphatic organochlorines were detected in sediment traps from the Arabian Sea, and their origin was suggested to be *G. huxleyi* following *in*-*situ* and culture-based measurements (33). These findings highlight the relevance of the still largely unknown biotic sources for organochlorines in the surface ocean. Finding aliphatic chloro-iodo metabolites in several virus-infected *G. huxleyi* blooms in the Atlantic Ocean suggests that alga-virus interactions are an important biotic source for organohalogens in the marine environment.

The chloro-iodo metabolites are likely persistent metabolic biomarkers in the marine environment owing to their polyhalogenation with up to four halogens. Culture-based studies showed that the majority of cyanobacterial vDOM (80%) gets respired within 48 hours (8). However, chloro-iodo metabolites require multiple dehalogenations before their metabolic degradation. This may explain their rapid accumulation during the mesocosm experiment and stable occurrence over several days during the Lagrangian drifts with the North Atlantic blooms. In turn, this may further lead to their sequestration as recalcitrant DOM without effective dehalogenation processes. Deiodination can occur by photolysis due to the low C-I bond energy (30) and depends on solar radiation and water circulation. Boreal *G. huxleyi* blooms, with their proximity to areas of deep-water formation, are thus more likely to lead to the transport of chloro-iodo metabolites into the deep sea and sequestration as recalcitrant DOM. Dechlorination is rather driven by microbial dehalogenation due to the photostability of the C-Cl bond. Polyhalogenated compounds, such as the chloro-iodo metabolites, are more persistent to microbial dehalogenation, which facilitates their transport and accumulation in the DOM (34) and POM (33) pools of the deep sea. Future *in*-*situ* experiments using on-deck incubations and sediment traps will help to shed light on photolytic and microbial dehalogenation processes and to model the transport and sequestration of chloro-iodo metabolites in the marine carbon cycle.

For a comprehensive understanding of the consequences of viral infections at the ecosystem level, biomarkers for other infection outcomes need to be developed, including the trophic transfer of carbon through selective feeding on virocells, or carbon export to the ocean floor through the viral shuttle (35). The development of such biomarkers is particularly relevant considering the global climate change affecting carbon cycling in the marine environment in unprecedented ways. Accordingly, there is a critical need for diagnostic tools that inform about the impact of climate change on the abundance, activity, and health of marine phytoplankton communities. For example, satellite remote sensing of surface water Chl *a* revealed changes in phytoplankton distribution and biomass (36) supported by *in*-*situ* observations that revealed further effects on biodiversity and species turnover (37). In addition, proxies are emerging that offer insights into phytoplankton activity, such as bio-optical measurements of primary production (38), which is a key trait of marine phytoplankton communities. Among the existing molecular biomarkers that can report on phytoplankton health are enzymes and lipids that indicate macro- and micronutrient limitations (39) and cell quotas in energy-rich lipids (40) and essential fatty acids (41) that have cascading nutritional effects at higher trophic levels. However, we still need biomarkers to monitor the prevalence and consequences of phytoplankton diseases, such as viral, bacterial, and fungal infections, which are rather understudied compared to marine macroorganisms. Environmental metabolomics, as applied in this study, proved to be crucial for finding ecologically relevant biomarkers, and will benefit from current efforts in implementing metabolite analyses in global ocean surveys.

## Methods

### Sampling and in-situ enumeration of G. huxleyi blooms

A summary of all samples with their relevant metadata is provided in Dataset S2.

#### AQUACOSM VIMS-Ehux mesocosm experiment

Blooms of *G. huxleyi* were induced by nutrient addition in four mesocosm bags at the Marine Biological Station Espegrend, Norway, between 23 May and 16 June 2018, as previously described (6). In brief, 40 µm pre-filtered water samples were analyzed immediately after sampling using an Eclipse iCyt flow cytometer (Sony Biotechnology, USA). Samples of 0.3 mL were run at 0.15 mL/min and calcified *G. huxleyi* cells were counted by plotting chlorophyll autofluorescence against side scatter.

#### NA-VICE cruise

Sampling of various *G. huxleyi* blooms along a transect in the North Atlantic was performed on board the R/V Knorr between 13 June and 16 July 2012, as described previously (20). A Lagrangian drift was performed using a sediment trap. Immediately after CTD sampling, unfiltered water samples were analyzed using a Guava flow cytometer (Millipore). Calcified *G. huxleyi* cells were enumerated by gating forward scatter against side scatter and chlorophyll fluorescence.

#### TARA Breizh Bloom cruise

Sampling of a *G. huxleyi* bloom patch in the Celtic Sea (North Atlantic) was performed on board the R/V Tara between 27 May and 2 June 2019, as described previously (42, 43). A Lagrangian drift was performed using an Argo float. For *G. huxleyi* enumeration, unfiltered water samples from a Niskin bottle were analyzed onboard immediately after sampling using a BD Accuri C6 Plus Personal Flow Cytometer (BD Biosciences, USA). Samples were run for 2 min at 66-69 µL/min. Calcified *G. huxleyi* cells were enumerated by plotting the chlorophyll fluorescence (ex.: 488 nm; em.: 663-737 nm) against the side scatter and gating the high-chlorophyll high-side scatter events.

#### TARA Ana María Gayoso cruise

Sampling of various *G. huxleyi* blooms along the Patagonian Shelf break (South Atlantic) was performed on board the R/V Tara between 4 and 27 December 2021 as part of a joint expedition with the R/V Houssay in the Mission Microbiomes AtlantEco project. The sampling in December targeted coccolithophore blooms, while the sampling in November on R/V Houssay targeted the pre-bloom conditions (Ferronato et al. *submitted*, Gilabert et al. *in press*). Lagrangian drifts were performed using SVP drifters. At each station, surface to maximum 100 m depth profiles of temperature, salinity, chlorophyll fluorescence and turbidity were conducted by deploying a SBE911+ profiler (Sea-Bird Scientific). Water samples were collected approximately every 10 m using Niskin bottles of 8 and 12 L. The maximum bloom depth was defined as the maximum *G. huxleyi* cell abundance determined using flow cytometry. Immediately after collection, 70 µm pre-filtered water samples were analyzed using a BD Accuri C6 Plus Personal Flow Cytometer (BD Biosciences, USA). Samples were run for 3 min at ∼60 µL/min. Calcified *G. huxleyi* cells were enumerated by gating chlorophyll fluorescence against side scatter as described above. Subsequent casts collected water for metabolite profiling and DNA extraction at the maximum bloom depth following adjustment to the CTD profile.

### Satellite remote sensing of particulate inorganic carbon (PIC) and chlorophyll a (Chl a)

The spatiotemporal distribution of surface water PIC and Chl *a* was mapped for each bloom using the Level 3 Aqua-MODIS satellite daily data. To generate the composite images of PIC visualized in Figure 1, daily maps were averaged between 23-26 June 2012 (V1-V2), 29 May and 1 June 2019 (B1_a-e_), 07-09 December 2021 (G1-G2), 15-18 December 2021 (G3_a-d_), 12-14 December 2021 (G51_a-c_) and 23-24 December 2021 (G4 and G6). For the composite images of PIC, Chl *a* and their ratio shown in Supplementary Figure S1, daily maps were averaged according to the time range indicated next to each image. The temporal profiles of Chl *a* and PIC of the Breizh bloom (Figure S1c) were generated by averaging 0.0417° (∼86 km^2^) around the sampling locations (48.3- 48.4°N, 6.9-7.1°W; outlined by white square), if cloud coverage permitted. For each sampling location, the respective surface water PIC, Chl *a* and Chl *a*:PIC ratio at the day of sampling are given in Supplementary Table S1. Based on the temporal dynamics of Chl *a*, PIC and the Chl *a*:PIC ratio, the bloom phase was defined following a previous conceptual model (20). In short, coccolithophore blooms are thought to initiate with an increase in PIC and, to a lesser degree, Chl *a* during the bloom phase, resulting in a parallel decrease in the Chl *a*:PIC ratio. During the onset of bloom demise, Chl *a* rapidly drops while PIC remains high due to the shedding of coccoliths from dead cells, resulting in a minimum Chl *a*:PIC ratio. Late bloom demise and phytoplankton succession will lead to an attenuation of PIC, an increase in Chl *a*, and thus a recovery of the Chl *a*:PIC ratio.

### Enumeration of extra- and intracellular EhV

To quantify extracellular and intracellular abundances of the *Emiliania huxleyi* virus (EhV), water samples were filtered, DNA extracted, and viral abundance analyzed by qPCR using specific primers for the major capsid protein (*mcp*) gene of EhV. The *mcp* primers designed by Pagarete and Schroeder were used for the NA-VICE cruise (22), while for all other samples, modified primers were used (44). For the <u>mesocosm experiment</u>, DNA was extracted from 0.2 µm and 2 µm filters (45). For the <u>NA-VICE cruise</u>, DNA was extracted from 0.8 µm filters as described in (20) (station V1) and (22) (station V2). For the <u>Breizh cruise</u>, DNA was extracted from 3 µm filters and analyzed by ddPCR (45). For the <u>Gayoso cruise</u>, up to 20 L (max. 15 min) of 20 μm pre-filtered water samples were filtered onto 3 μm polycarbonate filters (142 mm in diameter; Millipore). Filters were flash-frozen in liquid nitrogen and stored at -80°C until DNA extraction (46). qPCR analysis was performed as described previously (6) with slight modifications. All reactions were performed using the PowerUp SYBR Green Master Mix (Applied Biosystems) and the following program: 50°C for 2 min, 95°C for 2 min, and 40 cycles of 95°C for 15 s and 60°C for 60 s.

### Metabolite profiling of chloro-iodo vDOM metabolites across blooms

To screen for the chloro-iodo metabolites that characterize the vDOM of *G. huxleyi*, filter and filtrate samples of the induced mesocosm blooms and oceanic blooms were analyzed as described previously for the mesocosm experiment (6) with the following modifications.

Briefly, for the <u>mesocosm experiment</u>, 1-6 L of water were collected with a peristaltic pump through a 200 µm nylon mesh and then filtered sequentially through pre-combusted GF/A filters (47 mm; Whatman) and pre-cleaned 0.22 µm PVDF filters (47 mm; Durapore) by gentle vacuum filtration. Dissolved metabolites were extracted from 1 L aliquots of 0.22 µm filtrates that were acidified to pH 2.0.

Briefly, for the <u>VICE cruise</u>, 1-1.5 L of water were collected with a CTD rosette sampler, pre-filtered with a 200 µm mesh and then filtered through 0.8 µm PC filters (47 mm; Millipore). Particulate metabolites were extracted with methanol.

For the <u>Breizh cruise</u>, 50 L of water from the maximum bloom depth of *G. huxleyi* were pumped through a 20 µm nylon net and then filtered sequentially through pre-combusted GF/C filters (125 mm; Whatman) and 0.45 µm PVDF filters (142 mm; Durapore) using a vacuum pump. Filters were flash-frozen and stored in liquid nitrogen and -80°C. An aliquot of 1 L of the 0.45 µm filtrate was collected in a glass bottle and loaded with a peristaltic pump onto solid-phase extraction (SPE) cartridges (Oasis HLB, 500 mg; Waters) without prior sample acidification and SPE conditioning. Cartridges were flash-frozen in liquid nitrogen and stored at -20°C and 4°C until further processing seven months later.

For dissolved metabolite analysis, the cartridges were thawed, excess water removed by vacuum, and the cartridges washed (18 mL water; ULC/MS grade, Biolab), dried by vacuum, and eluted with 4 mL methanol (HPLC grade, J.T.Baker). For metabolomics analysis, eluates were dried under nitrogen flow, and re-dissolved in 200 μL methanol:water (1:1, v/v; ULC/MS grade water, BioLab) containing caffeine-(trimethyl-d_9_) (1.5 μg mL^-1^; 98%, Sigma-Aldrich) as injection standard. 110 μL were transferred to injection vials of which 1 µL was injected in positive and negative ionization modes for LC-MS analysis. The sample with the highest metabolite intensities (B2) was injected at 5 μL injection volume for MS/MS analyses in negative ionization mode. Peak areas were integrated using QuanLynx, and areas above an S/N of 6 were normalized to the injection standard (positive ionization mode) or TIC (negative ionization mode).

For the <u>Gayoso cruise</u>, 50 L of water were collected from the maximum bloom depth of *G. huxleyi* using a CTD rosette sampler, pumped through a 20 µm nylon net, and then filtered sequentially on pre-combusted GF/A filters (142 mm; Whatman) and pre-cleaned 0.22 μm PVDF filters (142 mm; Millipore) using a peristaltic pump. Aliquots of 1 L of filtrate were collected in glass bottles and loaded with a peristaltic pump onto SPE cartridges (Oasis HLB, 500 mg; Waters) following the addition of 5 µL d_2_-indole-3-acetic acid (d_2_-IAA, 1 μg/μL in water; ≥98%, Santa Cruz Biotechnology) as extraction standard. For the conditioning and equilibration of the cartridges onboard, LC-MS grade water and methanol (Merck LiChrosolv) were used. After cartridges were washed and excess water removed, samples were stored at -20°C until further processing seven months later.

After thawing, cartridges were washed with 18 mL water (Bio-Lab ULC-MS grade), dried, and eluted with methanol. For subsequent LC-MS analysis, dried eluates were re-dissolved in 120 μL methanol:water (1:1, *v:v*) containing caffeine-(trimethyl-d_9_) (1.5 μg mL^-1^) and d_5_-tryptophan (2.1 µg mL^-1^; 98%, Cambridge Isotope Laboratories) as injection standards. From supernatants, 110 μL were transferred to glass inserts for LC-MS analysis, and the remaining 10 μL combined in a pooled quality control (QC) sample. A blank sample was prepared at station G4 by processing artificial seawater along with the biological samples. Of each sample, 1 µL was injected in positive and negative ionization modes. The sample with the highest metabolite intensities (G2) was subjected to MS/MS analyses in positive and negative ionization modes, using an injection volume of 3 μL and 5 μL, respectively. Peak areas were integrated using QuanLynx and areas above an S/N of 6 were normalized to the injection standard d_5_-tryptophan.

All samples were screened for the presence of 19 vDOM chloro-iodo metabolites (#3 to 8, 10, 14, 16, and 21 to 30; see Dataset S2). Metabolites were initially identified based on the retention time and presence of indicative adduct ions of the M, M+2, and M+4 isotopes. Identities were further confirmed by the presence of the indicative fragment ion of iodine in MS/MS spectra in negative ionization mode. Peak areas were integrated for indicative adducts ([M+H]^+^, [M+H-H_2_O]^+^, [M-H]^-^) using QuanLynx. Peak areas above a certain S/N ratio were normalized depending on the sampling campaign as described above. To map the metabolite occurrence across blooms, metabolites were grouped by the halogenation level (from mono- to trichlorinated metabolites) and manually sorted by the frequency of their occurrence across all samples. A balloon plot was generated using the R package ‘ggpubr’. The size of the circles was scaled from 0 to 1 for each sampling campaign.

### Comparison of intra- and extracellular pools of chloro-iodo metabolites

To compare intra- and extracellular pool of chloro-iodo metabolites during the induced mesocosm blooms, which were sampled daily over 23 days, their concentrations in filter (1.6-25 µm) and acidified filtrate (<0.22 µm) samples from the strongly infected bag 4 were quantified.

The dissolved metabolite concentrations were analyzed as described previously (6).

For the extraction of particulate metabolites, 25 µm pre-filtered water samples were filtered gently through pre-combusted GF/A filters and spiked with 5 μL of internal standards before filters were flash-frozen in liquid nitrogen and stored at -80°C until further processing. Filters were lyophilized within 1.5 months after collection and then processed in four randomized batches. After thawing, filters were extracted twice by adding 3 mL pre-cooled (-20°C) methanol (HPLC grade, J.T. Baker) followed by 1 min vortexing, 30 min orbital shaking (darkness and 4°C) and 30 min sonication (100%, 37 Hz, <14°C). Extracts were then centrifuged (5 min, 3200×g, 4°C) and supernatants transferred to glass vials that were kept at 4°C during the 2^nd^ extraction of the filters. Supernatants were dried under a nitrogen flow and stored at -80°C until further processing. To remove filter debris, dried extracts were re-dissolved in 1.4 mL pre-cooled methanol, vortexed for 1 min, sonicated for 10 min and centrifuged for 10 min at 20,800×g. Supernatants were dried under a nitrogen flow and stored at -80°C until LC-MS analysis. A pooled QC sample was generated by combining 50 µL aliquots of each biological sample. Two filter blanks from the 4th and 18^th^ day of the mesocosm experiment were processed along with extraction solution blanks for each batch.

For LC-MS analysis, samples were re-dissolved in 200 µL methanol:water (1:1, v:v) and aliquots of 2 µL injected and analyzed in a comparative untargeted metabolite profiling approach as described previously for the dissolved metabolite extracts (6).

All samples were screened for the chloro-iodo metabolites using QuanLynx. Peak areas were integrated for the [M+H]^+^ or [M+H-H_2_O]^+^ adduct ions in positive ionization mode (#3-16) and or [M-H]^-^ adduct ion in negative ionization mode (#21-30) above a signal-to-noise (S/N) threshold of 10. Areas were normalized to the extraction standard d_2_-IAA and, for the filter samples, the filtration volume. As no internal standards were added on days 1-2 of the mesocosm experiment, the respective value of day 3 of each bag was used for normalization.

### Temporal profiling of different halogen-containing dissolved metabolites

To assess the general occurrence and temporal dynamics of chlorinated, brominated and iodinated metabolites throughout the induced mesocosm blooms of *G. huxleyi*, an UHPLC-HR-MS-based untargeted metabolite profiling (47) was performed for two types of pooled QC samples, one pooling days 1-23 and one pooling days 18-23 of the mesocosm bags 1-4. An aliquot of 1 µL was analyzed in alternating positive and negative ionization modes at 70,000 mass resolution with 100 ms maximum ion time. Isotopic pattern search for Cl, Cl_2_, Br and Br_2_ was run using the Pattern Scoring of Compound Discoverer (version 3.3, Thermo Fisher Scientific), following retention time alignment, peak picking and deconvolution. Compounds with predicted elemental compositions were manually validated and re-assigned within the full mesocosm metabolomics dataset (6) using their isotope pattern and retention time, and the chloro-iodo-containing metabolites among the chlorinated metabolites were labeled. A target mass list was compiled based on *m*/*z* and retention time. Peak areas of all target masses above an S/N threshold of 10 were integrated from days 10- 23 for bags 1-4 and the fjord using QuanLynx as described previously (6), normalized by the extraction standard d_2_-IAA and rescaled between 0-1. Two compounds that were always more abundant in the fjord were excluded from further analysis. For the remaining 5 chlorine-containing, 6 bromine-containing and 3 chloro-iodo-containing metabolites, the normalized peak areas were averaged per day and plotted using the ‘ggplot’ package.

### Correlation of chloro-iodo metabolites with environmental variables

To correlate the abundance of chloro-iodo metabolites with abiotic and biotic environmental parameters, a Spearman’s rank correlation analysis was performed for the mesocosm dataset. For each bag during the bloom and demise phases (days 10-23), the sum of all chloro-iodo metabolites was calculated both for the intra- and extracellular organic matter pool. Environmental variables included several core variables reported previously (45, 48), namely flow cytometry counts of *G. huxleyi*, nano- and picoeukaryotes, *Synechococcus* and bacterial cells, intra- and extracellular EhV abundances, chlorophyll *a*, as well as water temperature, salinity, and nutrient concentrations (nitrate and phosphate). GSLs were measured as described in (23), and the sum of all vGSLs was calculated per bag and day. Viral infection-induced transparent exopolymer particles (TEP) were quantified as described in (45), while algal DMSP and DMS were analyzed as described in (49). Zeros were treated as missing values to avoid data censoring due to analytical thresholds. Spearman’s rank correlation coefficients and 95% confidence intervals (CIs) were calculated using the R package ‘correlation’ and visualized as Forest plot using the ‘ggplot’ package. A correlogram of the Spearman’s correlation coefficient matrix was plotted using the R package ‘corrplot’. Parameters were ordered by their contribution to the first principal component that explains most of the variance in the dataset. There was no significant autocorrelation of the total abundance of chloro-iodo metabolites in each bag and the data was thus treated as independent measurements.

### Comparison of intracellular infection markers and vDOM markers in an open ocean bloom

To compare the presence of vGSLs as intracellular infection markers and chloro-iodo metabolites as vDOM markers in an oceanic *G. huxleyi* bloom, a lipidomics and metabolomics analysis was performed for filter (1.2-20 µm) and non-acidified filtrate (<0.45 µm) samples of the intense Breizh bloom over a 5-days Lagrangian drift during demise phase.

Dissolved metabolite analysis and screening for the chloro-iodo metabolites was performed as described above.

For the extraction of particulate metabolites, filters were lyophilized and then extracted and analyzed as described previously (23) with slight modifications. Filters were extracted using 20 mL methanol:MTBE (1:3, v:v) containing Cer/Sph internal standard mixture I (22.5 nM; Avanti) and 15 mL water:methanol (3:1, v:v) for phase separation. Re-extraction of the filters was performed with 11 mL MTBE and 6 mL methanol for lipophilic and polar metabolites, respectively. In total, 26 mL upper organic phase was collected for lipidomics analysis and 15 mL lower polar phase for metabolomics analysis. A filter blank and extraction solution blank were processed along with the biological samples.

For lipidomics analysis of the GSLs, filter extracts were dried under nitrogen flow, re-dissolved in 300 µL re-dissolving solution, and 200 µL of the supernatant transferred to injection vials as described previously (23). Prior to injection, samples were diluted threefold and 1 µL analyzed in positive ionization mode. Identification of GSL species was based on fragments in MS^E^ mode and peak areas integrated for indicative adduct and fragment ions following collision-induced dissociation in MS^E^ mode ([M+Na]^+^ for hGSLs and vGSLs, and [M+H-(Sialic acid-H)-H_2_O]^+^ for sGSLs) using QuanLynx (0.05 Da mass window). Peak areas above an S/N threshold of 10 were normalized to the extraction standard glucosyl(beta)C12 ceramide.

For metabolomics analysis of the chloro-iodo metabolites, filter extracts were dried under nitrogen flow, re-dissolved in 300 µL methanol:water (1:1, v:v) with caffeine-(trimethyl-d_9_)(1.5 μg mL^-1^) as injection standard, centrifuged, and 220 µL of the supernatant transferred to injection vials. For LC-MS analysis, 2 µL were injected in positive ionization mode. Peak areas were integrated using QuanLynx, and areas above an S/N of 10 were normalized to the injection standard.

## Data availability

All data needed to evaluate the conclusions in the paper are present in the paper, the Supporting Information, and Dataset S1-S2.

## Acknowledgments

We thank all team members of the AQUACOSM VIMS-Ehux project for setting up and conducting the mesocosm experiment, especially Jorun Egge, Aud Larsen and Tatiana Tsagaraki. We further thank the chief scientist of the NA-VICE cruise, Kay Bidle, the captain and crew of the R/V Knorr, and the Marine Facilities and Operations at the Woods Hole Oceanographic Institution for their assistance and cooperation. We wish to thank the Tara Ocean Foundation, the SV Tara crew, notably captain Samuel Audrain and Martin Herteau, and Colomban de Vargas for the sampling opportunity and scientific and logistic support during the Breizh and Gayoso cruises. Furthermore, we wish to thank all participants of Mission Microbiomes AtlantECO, particularly Daniele Iudicone and Stephane Pesant, and the Argentinian authorities and scientists, especially Pedro Flombaum, Valeria Guinder, Federico Ibarbalz, Celeste López-Abbate and Martin Saraceno, for their support during the Gayoso cruise. We thank the French National Sequencing Center (CEA-Genoscope, Evry) for sending aliquots of DNA extracts for qPCR analysis. We also thank the commitment of the following institutions for their financial and scientific support that made Mission Microbiomes AtlantECO possible: Stazione Zoologica Anton Dohrn, European Bioinformatics Institute (EMBL-EBI), Centre national de la recherche scientifique (CNRS), Centre National de Séquençage (CNS, Genoscope), agnès b., BIC, Capgemini Engineering, Fondation Groupe EDF, Compagnie Nationale du Rhône, L’Oréal, Biotherm, Région Bretagne, Lorient Agglomération, Billerudkorsnas, Havas Paris, Fondation Rothschild, Office Français de la Biodiversité, AmerisourceBergen, Philgood Foundation, UNESCO-IOC, Etienne Bourgois. This work was supported by a European Research Council Advanced Grant (no. 101053543; VIBES), a Simons Foundation grant (no. 735079; ‘Untangling the infection outcome of host-virus dynamics in algal blooms in the ocean’), and a Minerva-Weizmann Project Grant (no. 140686) awarded to A.V. The mesocosm experiment VIMS-Ehux was supported by EU Horizon2020-INFRAIA project AQUACOSM (no. 731065), and the NA-VICE cruise was supported by an NSF grant (no. OCE- 1059884). MV is supported by the Deutsche Forschungsgemeinschaft (DFG, German Research Foundation) project SFB1127 ChemBioSys (no. 239748522).

## Competing Interest Statement

The authors declare no competing interest.

## Author Contributions

C.K. and A.V. conceptualized the study and experimental design and wrote the manuscript. C.K. and G.S. performed the metabolomics experiments. J.M.F. processed and analyzed the satellite data. F.V. and D.S. performed flow cytometry analyses during TARA Breizh and Gayoso. C.K., D.S., G.S. and F.V. performed qPCR and ddPCR analyses for TARA Breizh and TARA Gayoso. M.V. generated mass spectral data that was used for the isotope pattern search. All authors reviewed and edited the manuscript.

## Notes

### Competing Interest Statement

The authors have declared no competing interest.

