## Supporting Information for "Mapping of the viral shunt across widespread coccolithophore blooms using metabolic biomarkers"

**Author Contributions:** C.K. and A.V. conceptualized the study and experimental design and wrote the manuscript. C.K. and G.S. performed the metabolomics experiments. J.M.F. processed and analyzed the satellite data. F.V. and D.S. performed flow cytometry analyses during TARA Breizh and Gayoso. C.K., D.S., G.S. and F.V. performed qPCR and ddPCR analyses for TARA Breizh and TARA Gayoso. M.V. generated mass spectral data that was used for the isotope pattern search. All authors reviewed and edited the manuscript.

**Competing Interest Statement:** The authors declare no competing interest.

**Keywords:** algal bloom, *Gephyrocapsa huxleyi*, alga-virus interaction, virocell, vDOM, halogenation, biomarker

**This file includes:**

Figures S1 to S4
Legends for Datasets S1 to S2

**Other Supporting Information files:**

Datasets S1 to S2 (.csv)

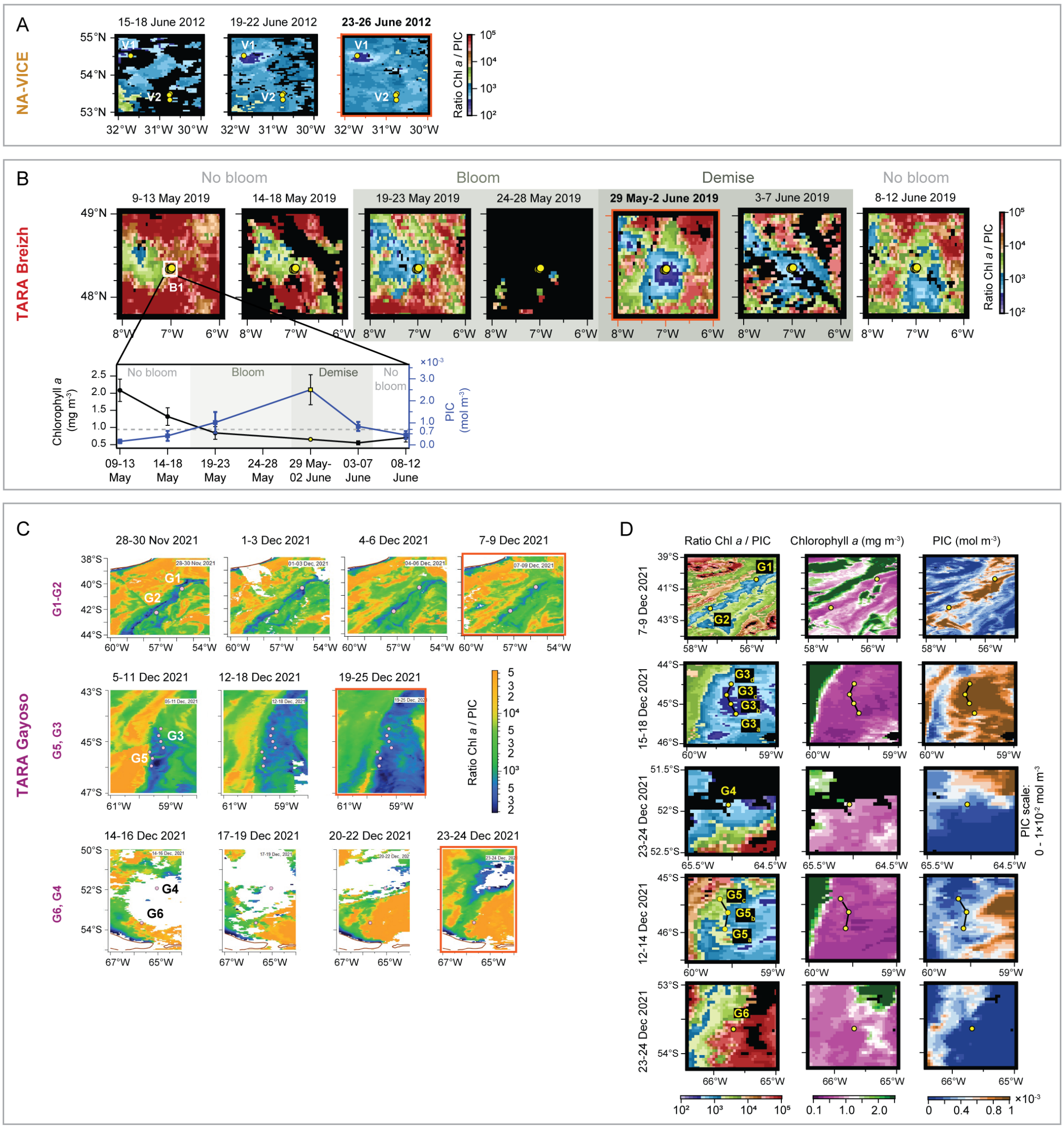


**Figure S1. Temporal dynamics of oceanic *G. huxleyi* blooms mapped by satellite image analysis of Chl *a*:PIC ratio, Chl *a*, and PIC.** Coccolithophore blooms are characterized by low Chl *a*:PIC ratios. A) Temporal dynamics before and during the NA-VICE bloom sampling. B) Temporal dynamics of Chl *a*:PIC before, during and after the TARA Breizh bloom sampling. White square (48.3-48.4°N, 6.9-7.1°W) marks the area that was averaged for the inset showing a time course of Chl *a* and PIC. C) Temporal dynamics of Chl *a*:PIC before and during the Gayoso bloom sampling in December 2021. D) Detailed maps of the Chl *a*:PIC ratio, Chl *a* and PIC. Circles indicate sampling locations of all cruises. Red plot outline highlights the maps that cover the sampling period.

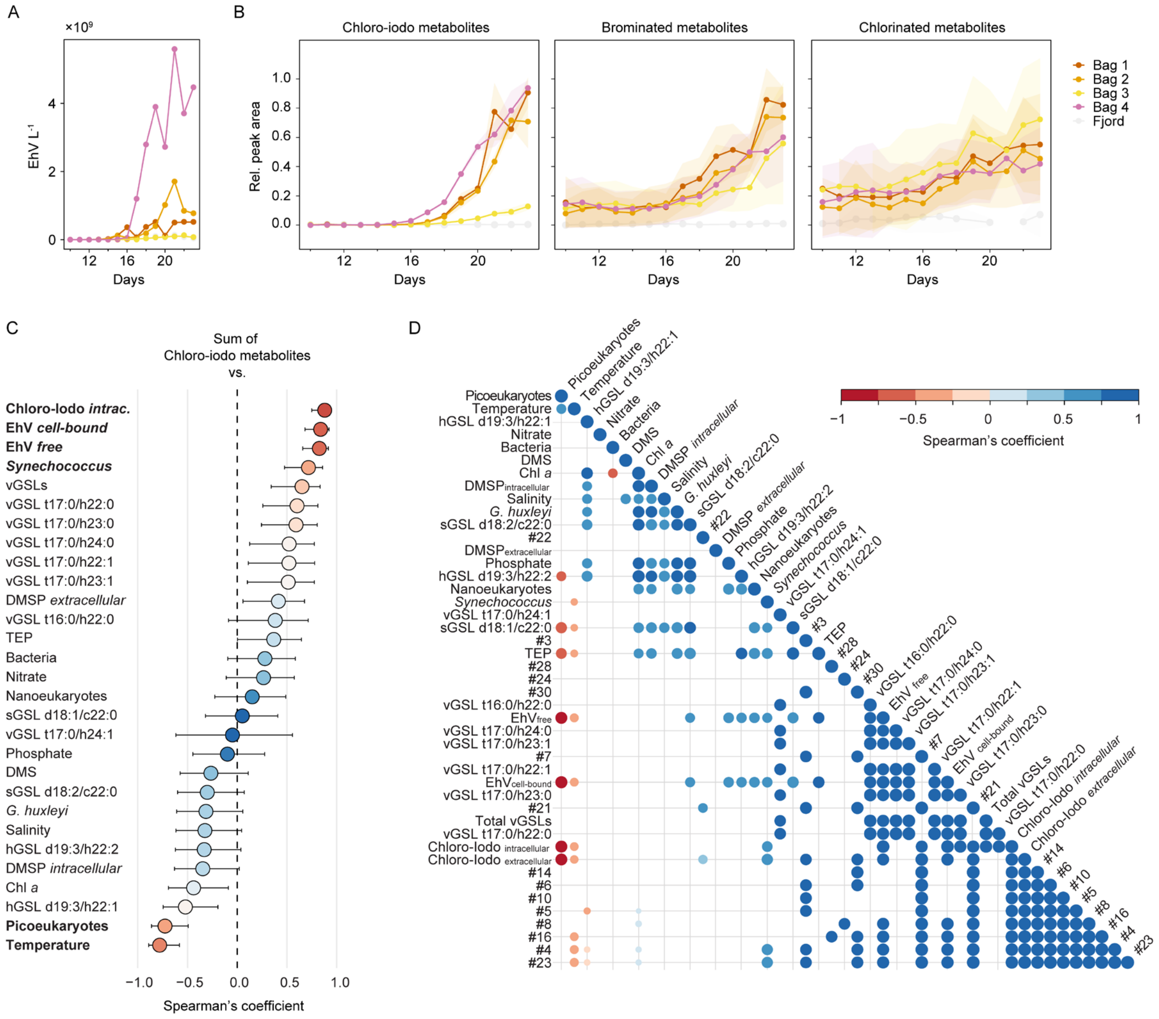


**Figure S2. Occurrence of other halogen-containing metabolites and correlation of chloro-iodo metabolites with diverse environmental parameters during induced mesocosm blooms.** A) Extracellular viral abundance throughout the bloom and demise phases of *G. huxleyi* in four mesocosm bags and the surrounding fjord showing their different level of infection. Quantification of EhV in seawater filtrates (<0.22 µm) was based on qPCR of a viral gene marker (*mcp* copies $L^{-1}$). B) Extracellular abundance of chloro-iodo metabolites (n=3) as compared to brominated (n=6) and chlorinated (n=5) metabolites. Average ± standard deviation of relative peak areas after normalization to the internal standard and scaling to the maximum intensity of each metabolite. The surrounding fjord served as an environmental reference. C) Spearman's rank correlation coefficient and 95% confidence interval (CI) for pairwise correlations of the total abundance of chloro-iodo metabolites with environmental parameters in mesocosm bags 1-4 during the bloom and demise phases (days 10-23). Environmental parameters include intracellular chloro-iodo metabolites, free and cell-bound EhV particles, virus-derived glycosphingolipids (vGSLs), general host-derived GSLs (hGSLs) and GSLs specific for virus-susceptible cells (sialic acid GSLs, sGSLs), cell counts of *G. huxleyi*, nano- and picoeukaryotes, *Synechococcus* and bacteria, chlorophyll *a* (Chl *a*), transparent exopolymeric particles (TEP), dimethylsulfide (DMS), intra- and extracellular dimethylsulfoniopropionate (DMSP), water temperature, salinity, nitrate and phosphate concentrations. Significantly correlated parameters are written in bold. D) Correlogram depicts significant pairwise correlations between each parameter. Extracellular concentrations of summed and individual chloro-iodo metabolites are presented. Parameters are ordered by their contribution to the first principal component.

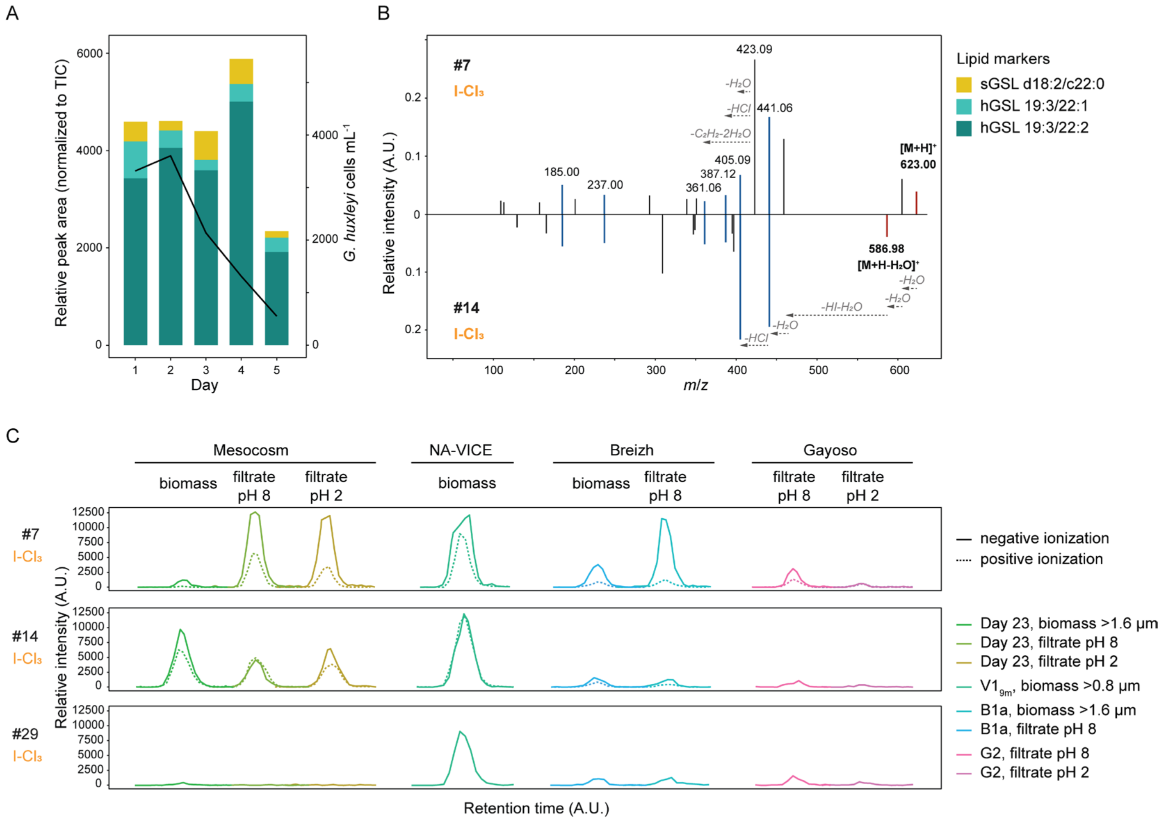


**Figure S3. Mass spectral characterization of trichloro-iodo metabolites that can serve as sensitive markers for the viral shunt in *G. huxleyi* blooms.** A) Temporal profiles of lipid infection markers in the biomass samples (1.2-20 µm) from a 5-days Lagrangian drift with the demising Breizh bloom (B1a-e). Lipids markers for all *G. huxleyi* cells (host-derived GSLs, hGSLs) and virus-susceptible cells (sialic acid GSLs, sGSLs) were found, while no markers of virus-infected cells were detected (virus-derived GSLs, vGSLs). Cell abundance of *G. huxleyi* (solid line) show a decrease in calcified *G. huxleyi* cells. Peak areas were normalized to the total ion current. B) MS/MS spectra of metabolites #7 (top) and #14 (bottom) in positive ionization mode share six major fragments (highlighted in pink), suggesting that these metabolites are structurally related. Their precursor ions (highlighted in yellow) differ by a loss of water ($H_2O$). Characteristic neutral losses are indicated. C) Extracted ion chromatograms of metabolites #7 (top), #14 (middle), and #29 (bottom) in negative (solid line) and positive (dashed line) ionization modes across various sample types and extraction methods, highlighting their robust detection in natural blooms of *G. huxleyi*. Data include biomass samples with two filter pore sizes and seawater filtrates with (pH 2) and without (pH 8) acidification before metabolite extraction. These metabolic biomarkers will be detected at higher signal intensities in extracts of non-acidified seawater filtrates analyzed in negative ionization mode.

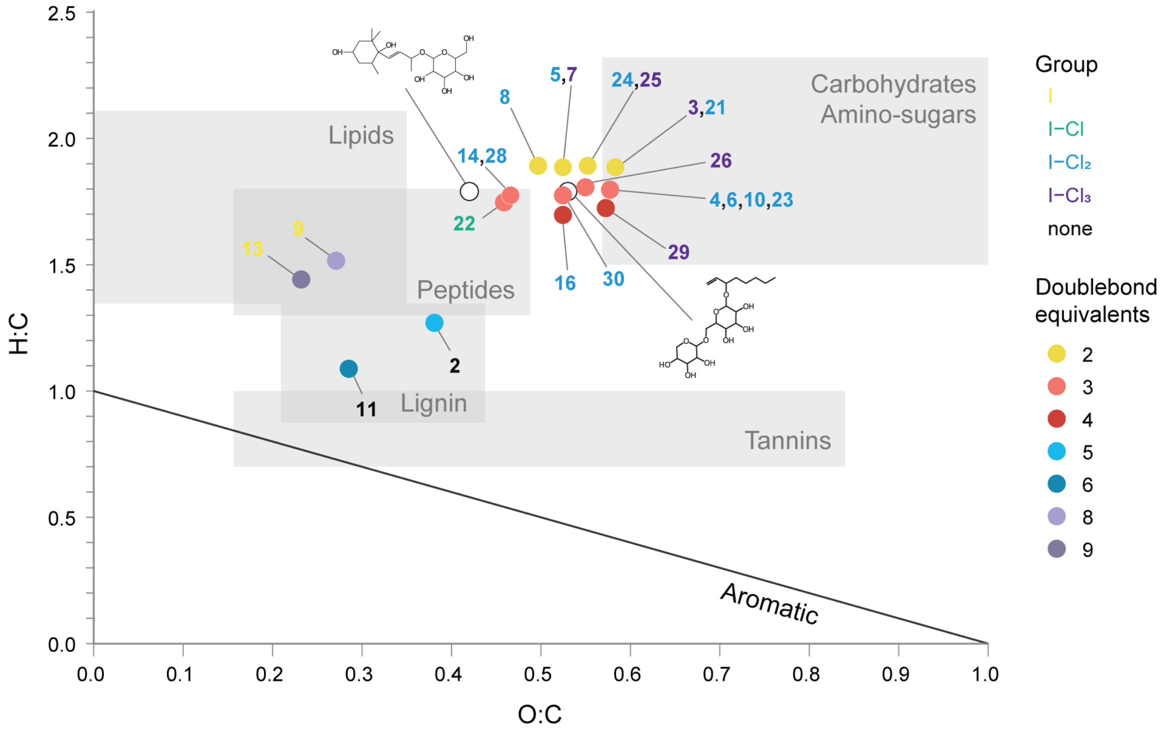


**Figure S4. Van Krevelen diagram of vDOM metabolites of *G. huxleyi*.** Ratios of H:C and O:C for the predicted elemental compositions of *G. huxleyi* vDOM metabolites. The label number represents the metabolite index, and the label color reflects the halogenation pattern regarding the number of iodine and chlorine atoms. The circle color reflects the number of double bond equivalents (DBEs) with one equivalent representing either a double bond or a ring. Grey rectangles outline the chemical space that known metabolites of common compound classes cover based on current chemical databases (13). The structures visualize known fatty acyl glycosides with stochiometric proportions comparable to the chloro-iodo metabolites (PubChem identifiers: left structure – 16745401, right structure – 24094121).

**Dataset S1. Metadata summary of all environmental samples.** Data sheet 1 includes cruise station data such as sample identifiers, sampling date, latitude, longitude, depth, satellite PIC and Chl *a*, flow cytometry cell counts, and qPCR data. Data sheet 2 includes mesocosm metadata such as physical and nutrient measurements, flow cytometry counts, biogeochemical analyses, viral abundances, lipid infection markers and chloro-iodo metabolite concentrations. (separate file, .csv)

**Dataset S2. Mass spectral data summary of all environmental samples.** Data sheet 1 includes metabolite identifier, halogenation type, retention time, m/z values and annotations, predicted elemental composition, van Krevelen parameters, number of double bond equivalents (DBEs), and normalized peak areas for each bloom sample. (separate file, .csv)
